# Advances in the integration of transcriptional regulatory information into genome-scale metabolic models

**DOI:** 10.1101/053520

**Authors:** R.P. Vivek-Ananth, Areejit Samal

## Abstract

A major goal of systems biology is to build predictive computational models of cellular metabolism. Availability of complete genome sequences and wealth of legacy biochemical information has led to the reconstruction of genome-scale metabolic networks in the last 15 years for several organisms across the three domains of life. Due to paucity of information on kinetic parameters associated with metabolic reactions, the constraint-based modelling approach, flux balance analysis (FBA), has proved to be a vital alternative to investigate the capabilities of reconstructed metabolic networks. In parallel, advent of high-throughput technologies has led to the generation of massive amounts of *omics* data on transcriptional regulation comprising mRNA transcript levels and genome-wide binding profile of transcriptional regulators. A frontier area in metabolic systems biology has been the development of methods to integrate the available transcriptional regulatory information into constraint-based models of reconstructed metabolic networks in order to increase the predictive capabilities of computational models and understand the regulation of cellular metabolism. Here, we review the existing methods to integrate transcriptional regulatory information into constraint-based models of metabolic networks.

## 1. Introduction

Extensive research by biochemists in the last century has resulted in the chemical characterization of thousands of biochemical reactions (Kanehisa and Goto, 2000; Chang et al., 2009). Towards the end of 20^th^ century, complete genome sequences became available for the first time (Fleischmann et al., 1995; Tomb et al., 1997). In the post-genomic era, a focus in systems biology has been the reconstruction of genome-scale metabolic networks using the annotated sequences along with available biochemical, genetic and phenotypic information for organisms (Edwards et al., 2001; Forster et al., 2003; Duarte et al., 2007; Feist et al., 2009; Thiele and Palsson, 2010; Thiele et al., 2013). In the last 15 years, considerable effort has led to the reconstruction of manually curated and high quality genome-scale metabolic networks for more than 50 organisms including humans (Durot et al., 2009; Feist et al., 2009; Thiele and Palsson, 2010). However, the current pace of manual reconstruction of high quality genome-scale metabolic networks lags far behind the sequencing effort, and thus, automated methods have also been developed to aid and accelerate the speed of metabolic network reconstruction process (Henry et al., 2010; Schellenberger et al., 2011; Agren et al., 2013). Several studies have demonstrated the utility of these genome-scale metabolic reconstructions for biological discovery and hypothesis generation (Feist and Palsson, 2008; Oberhardt et al., 2009).

Current paucity of information on relevant parameters such as rate constants, enzyme concentrations and metabolite concentrations for most reactions renders kinetic modelling of genome-scale metabolic networks infeasible. In the face of inadequate kinetic information, the constraint-based modelling method, flux balance analysis (FBA), has proved to be a vital alternative to study the capabilities of genome-scale metabolic networks (Varma and Palsson, 1994; Kauffman et al., 2003; Price et al., 2004; Orth et al., 2010; Lewis et al., 2012). In contrast to kinetic models, constraint-based FBA primarily uses stoichiometric information (Heinrich and Schuster, 1996; Schilling et al., 1999; Palsson, 2006) of reactions in a metabolic network to predict the flux of reactions in steady state and biomass synthesis rate in a given environmental condition. Due to its simplicity, constraint-based FBA has become a popular framework to study genotype-phenotype relationships and predict the metabolic response to environmental and genetic perturbations using genome-scale metabolic reconstructions (Feist and Palsson, 2008; Papp et al., 2009; Lewis et al., 2012).

Concurrently, advances in post-genomic high-throughput data collection techniques has led to the generation of vast amounts of *omics* data. High-dimensional multi-omics data provides quantitative information on multitude of cellular components across diverse scales of organization. A major goal of systems biology is to turn the recent explosion of omics data into predictive holistic models of biological systems (Kitano, 2002; Joyce and Palsson, 2006). In this direction, contextualization of omics data within constraint-based FBA models of genome-scale metabolic reconstructions can lead to more accurate models. However, there are several challenges in integration of omics data stemming from the inherent experimental and biological noise in such datasets (Quackenbush, 2004). Nonetheless, several constraint-based methods have been developed to integrate experimental data, especially on transcriptional regulation and gene expression, within the FBA framework to build improved models (Åkesson et al., 2004; Covert et al., 2004; Becker and Palsson, 2008; Blazier and Papin, 2012; Hyduke et al., 2013). In this review, we discuss the existing methods to integrate regulatory information into constraint-based FBA models by broadly classifying them into three different approaches (Covert et al., 2001; Åkesson et al., 2004; Covert et al., 2004; Becker and Palsson, 2008; Chandrasekaran and Price, 2010; Blazier and Papin, 2012; Hyduke et al., 2013; Kim and Reed, 2014). Such methods have already proven successful in building context-specific metabolic models for human tissues and predicting novel drug targets in pathogens (Becker and Palsson, 2008; Folger et al., 2011; Bordbar et al., 2012; Collins et al., 2012).

The review is organized as follows. In the second section, we describe the constraint-based FBA framework. In the third section, we discuss existing methods to integrate omics data within the FBA framework as additional flux constraints to build context-specific metabolic models. In the fourth section, we describe the reconstruction and analysis of integrated regulatory-metabolic models where Boolean transcriptional regulatory networks (TRNs) are incorporated within the FBA framework. In the fifth section, we discuss the need for automated methods to integrate information on regulatory network architecture and expression measurements within metabolic networks to reconstruct integrated regulatory-metabolic models. Note that previous reviews in this area only emphasize on methods that are descriptive in nature which are presented in section 3 of this review. In comparison to previous reviews, we here provide a much more comprehensive overview of the area by also describing in detail the methods which are predictive rather than just descriptive in nature in sections 4 and 5 of this review.

## 2. Flux balance analysis

Flux balance analysis (FBA) is a constraint-based modelling approach that is widely used to investigate the capabilities of available genome-scale metabolic networks (Varma and Palsson, 1994; Kauffman et al., 2003; Price et al., 2004; Orth et al., 2010; Lewis et al., 2012). FBA primarily uses the information on the list of biochemical reactions in an organism along with the stoichiometric coefficients of involved metabolites to predict the fluxes of all reactions in the metabolic network. Such biochemical information is contained within available organism-specific genome-scale metabolic reconstructions. For any organism, the genome-scale metabolic reconstruction contains information on all known metabolic reactions and genes encoding enzymes catalysing different reactions in the network (Palsson, 2006) (Fig. 1A). Notably, genome-scale metabolic reconstructions for most organisms also include reactions for transport of metabolites across the cell boundary, and a pseudo-reaction capturing the production of biomass in terms of their precursor metabolites (Fig. 1A).

**Figure 1:**
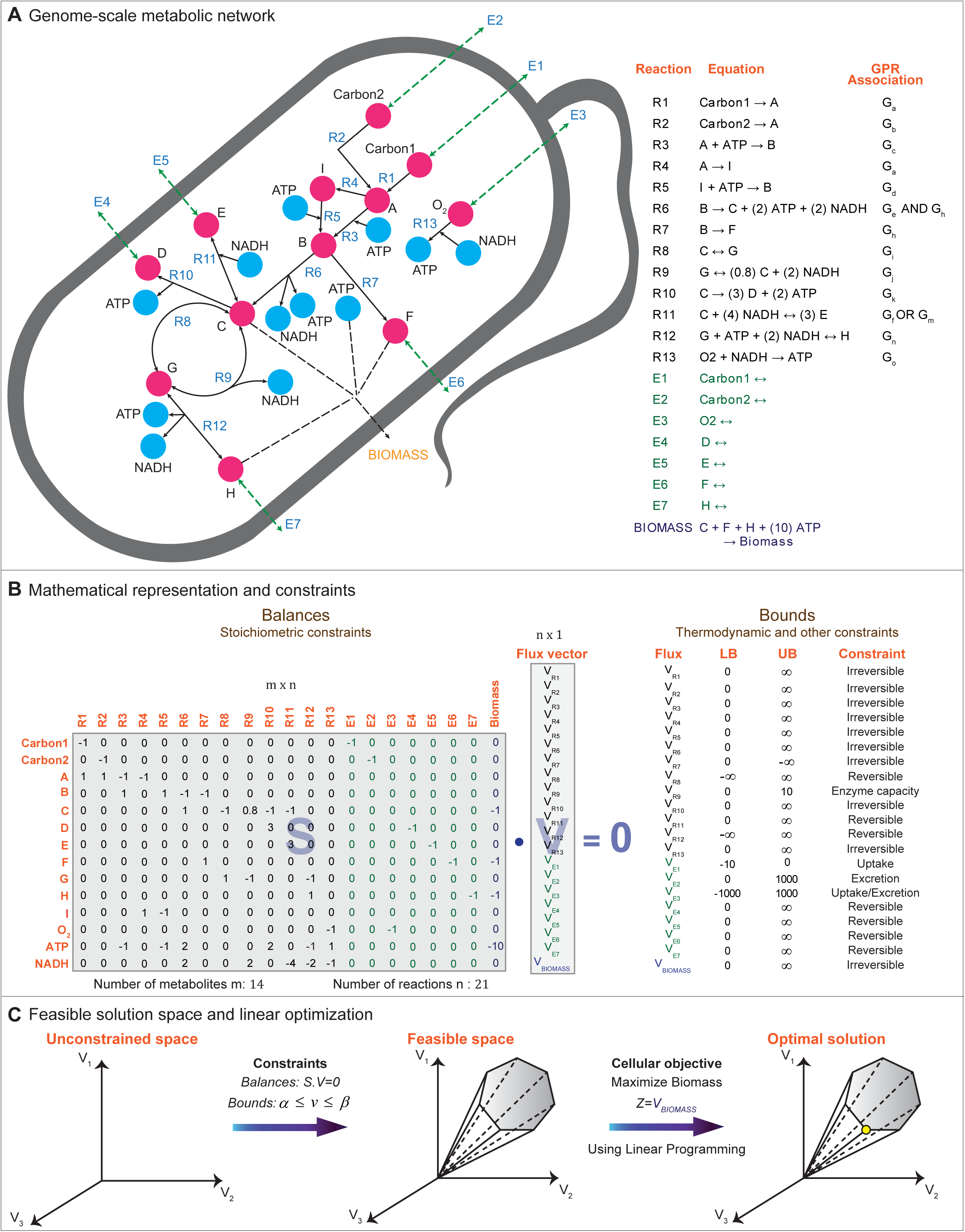
Overview of the FBA framework. **(a)** Genome-scale metabolic network reconstruction contains the list of metabolic reactions in an organism including the transport reactions and the biomass reaction. Gene-Protein-Reaction (GPR) associations link genes to encoded enzymes catalysing various reactions in the network, **(b)** The list of reactions along with the stoichiometric coefficients is mathematically represented in the form a matrix **S** where rows correspond to metabolites and columns to reactions in the network. In any metabolic steady state, the stoichiometric constraints lead to a system of mass-balance equations relating various reaction fluxes in the network. Additional constraints based on thermodynamic and other considerations bound the possible range of fluxes through specific reactions. **(c)** The stoichiometric and additional constraints lead to an under-determined system of equations with a large space of allowable solutions. FBA uses LP to find a particular solution within the allowable solution space that maximizes the biomass production. Here, the schematic figure shown in (c) is inspired from various sources including Orth *et al.* (Orth et al., 2010).

In the FBA framework, the list of reactions along with the stoichiometric coefficients of involved metabolites in a network reconstruction is mathematically represented in the form of a stoichiometric matrix **S** of dimensions *m* × *n*, where *m* denotes the number of metabolites and *n* denotes the number of reactions in the network (Fig. 1B). Entries in each column of the matrix **S** give the stoichiometric coefficients of metabolites participating in a particular reaction, where negative coefficients signify consumption of a metabolite, positive coefficients signify production of a metabolite, and zero coefficients signify no participation of a metabolite in the reaction (Fig. 1B). These stoichiometric coefficients of metabolites in various reactions impose constraints on the flow of metabolites in the network (Heinrich and Schuster, 1996; Schilling et al., 1999; Palsson, 2006). Subsequently, the method capitalizes on these stoichiometric constraints and assumes steady state to predict the fluxes of all reactions in the network.

In any metabolic steady state, different metabolites attain a mass balance wherein the rate of production of each metabolite is equal to its rate of consumption, and this leads to the system of mass balance equations given by:

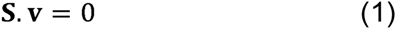
where **v** is the vector of fluxes through all reactions in the network (Fig. 1B). For each metabolite in the network, Eq. 1 gives a linear equation relating fluxes of various reactions in which the metabolite participates (Fig. 1B). Since, the number of metabolites is much less than the number of reactions in genome-scale metabolic networks of most organisms, the number of linear equations is much less than the number of reaction fluxes (unknowns) to be determined. Thus, Eq. 1 typically leads to an under-determined system of linear equations, and a large solution space of allowable fluxes for genome-scale metabolic networks (Fig. 1B,C).

The size of the allowable space can also be reduced by incorporating additional constraints on reaction fluxes. Firstly, certain reactions in the metabolic network are irreversible under physiological conditions, and such thermodynamic constraints (Beard et al., 2002; Orth et al., 2010) can be used to constrain the flux of irreversible reactions. Secondly, the activity of specific enzymes may limit the flux through certain reactions. Thirdly, the availability of nutrients in the growth medium can be used to constrain the fluxes of transport reactions. Note that unlike stoichiometric or mass-balance constraints, these additional constraints represent bounds on reaction fluxes in the metabolic network (Fig. 1B).

Since stoichiometric and additional constraints lead to an under-determined system with a large space of possible solutions, FBA uses linear programming (LP) to find a particular solution within the allowable solution space that either maximizes or minimizes a certain biologically relevant linear objective function *Z* (Watson, 1984; Fell and Small, 1986; Varma and Palsson, 1994) (Fig. 1C). Some examples of biologically relevant objective functions that have been explored using FBA include maximization of biomass production, maximization of ATP production and minimization of redox potential (Schuetz et al., 2007). However, maximization of biomass production (Feist and Palsson, 2010) is usually chosen as the objective function in FBA. The LP formulation of FBA can be written as:

Maximize *Z* = V_BIOMASS_ subject to

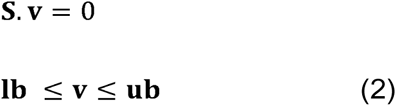
where v_BIOMASS_ is the objective function corresponding to the biomass growth flux, and vectors **lb** and **ub** contain the lower and upper bounds on different reaction fluxes contained in **v**. The solution of the FBA problem in Eq. 2 is a flux distribution **v** that maximizes the biomass objective function subject to the stoichiometric and additional constraints. FBA has found wide applications which include microbial strain improvement for producing industrially important metabolites, understanding molecular pathways involved in microbial pathogenesis, identification of drug targets, understanding reductive evolution in microbes and generating tissue-specific metabolic models for normal and cancerous tissues (Burgard et al., 2003; Pal et al., 2006; Becker and Palsson, 2008; Gianchandani et al., 2010; Folger et al., 2011; Bordbar et al., 2012; Collins et al., 2012).

## 3. Integration of transcriptomic profiles to build context-specific metabolic models

FBA predicts the fluxes of reactions and the biomass production rate without accounting for the regulatory constraints that are crucial in determining the presence and activity of enzymes in an environmental condition (Orth et al., 2010). This omission of regulatory constraints is an important limitation of FBA (Covert and Palsson, 2002, 2003; Åkesson et al., 2004) and can partly explain incorrect predictions by this method on gene essentiality (Covert et al., 2004) and gene interactions (Szappanos et al., 2011). The metabolic state of a cell in a given condition is governed by the expression of metabolic genes encoding enzymes. Thus, gene expression measurements obtained from microarrays or RNA-sequencing (RNA-seq) provide vital information on the state of the regulatory network and its influence on the metabolic network. With drastic reduction in cost of microarrays and RNA-seq, these high-throughput technologies are being increasingly used to generate abundant information on the expression of genes for several organisms under varied environmental conditions. Such data can be readily exploited to understand the condition-specific activity and regulation of metabolism. Towards this goal, several constraint-based methods have been developed in the last few years to integrate gene expression data within the FBA framework to generate context-specific metabolic models (Blazier and Papin, 2012; Lewis et al., 2012; Hyduke et al., 2013; Kim and Reed, 2014; Machado and Herrgård, 2014; Saha et al., 2014; Imam et al., 2015). The available methods can be broadly classified into three categories (Estévez and Nikoloski, 2014), namely, switch-and valve-based methods that use omics data to determine active and inactive set of genes in a given condition, methods that generate context-specific metabolic models without the need for a predefined biological objective function, and methods based on iterative removal of inferrred non-functional reactions to build tissue-specific metabolic models.

In the first category of methods, transcriptomic data is used to determine the set of active and inactive (absent) genes in a given condition (Fig. 2). Next, the information on active and inactive genes is used to set bounds on reaction fluxes catalysed by associated gene products (enzymes) within the FBA framework before computing the flux distribution and biomass rate. The treatment of bounds on reaction fluxes catalysed by active and inactive enzymes varies between different methods in this category, and this leads to a further classification into two sub-categories (Hyduke et al., 2013) (Fig. 2). In the sub-category of switch-based methods, the upper bound for the maximum flux through reactions catalysed by active enzymes is left unconstrained while those for reactions catalysed by inactive enzymes is set to zero before computing the flux distribution using FBA (Fig. 2). In the sub-category of valve-based methods, the upper bound for the maximum flux through reactions catalysed by enzymes is set proportional to the normalized expression of the associated genes before computing the flux distribution using FBA (Fig. 2). Basically, these methods are all based on the assumption that gene expression is correlated with reaction fluxes which may not necessarily hold (Gygi et al., 1999; ter Kuile and Westerhoff, 2001; Yang et al., 2002). Moreover, omics measurements suffer from a lack of sensitivity leading to false negatives in the identification of active and inactive genes. Notably, switching off flux through reactions catalysed by inactive genes which are false negatives due to experimental noise, may render the FBA predictions inconsistent with the biological objective or required metabolic function. Thus, the first category of methods also have the option to selectively re-enable flux through some reactions catalysed by inactive enzymes, to overcome any potential inconsistent FBA predictions with the required metabolic function (Becker and Palsson, 2008).

**Figure 2:**
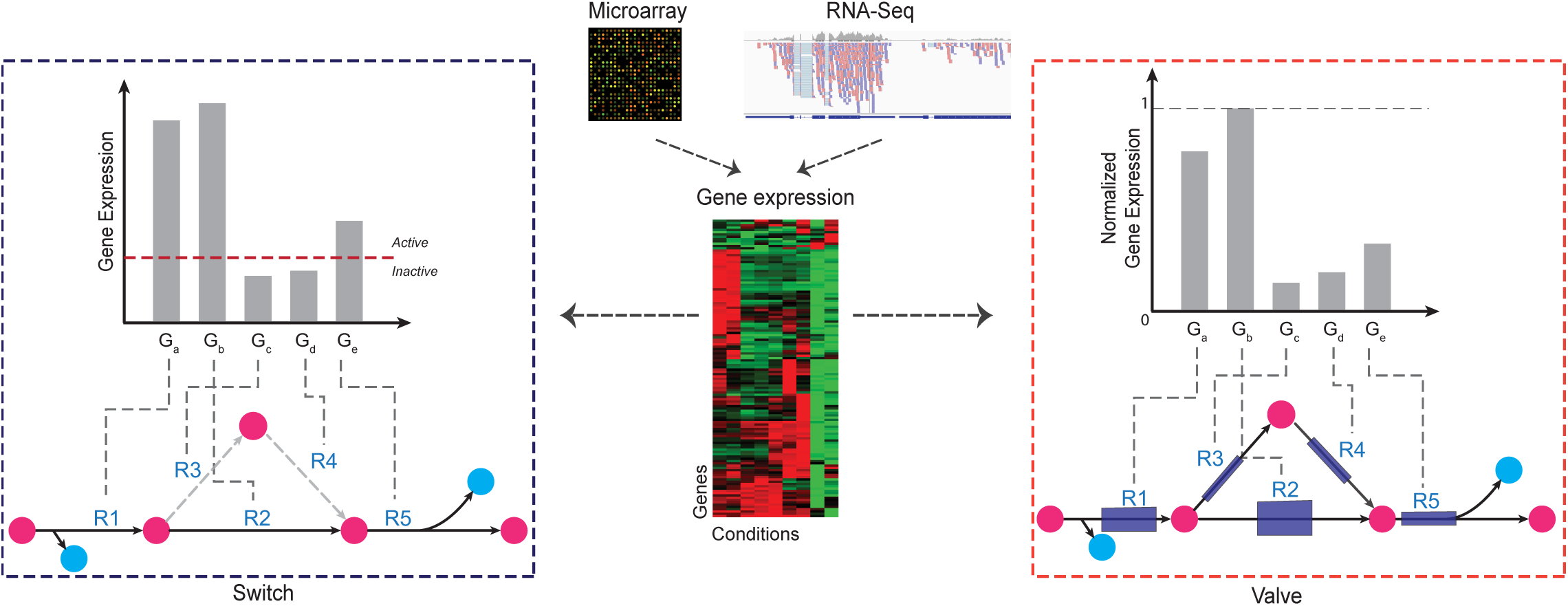
Integration of gene expression data within FBA framework using switch-and valve-based methods. Switch-based methods classify genes as active and inactive based on a threshold on expression value, and this leads to a binary classification of upper bounds on associated reaction fluxes. In the example network on the left, genes G_c_ and G_d_ are inactive, and thus, the upper bound on flux through their associated reactions R3 and R4 are set to zero (or, in other words, reactions R3 and R4 are switched off). In contrast, valve-based methods use normalized gene expression to set upper bounds on maximum allowable reaction fluxes. In the example network on the right, the magnitude of upper bounds on different reaction fluxes is represented via blue boxes with varying widths. Note that unlike in the switch-case, the upper bound on flux through reactions R3 and R4 are not set to zero in the valve-case.

Akesson *et al.* (Åkesson et al., 2004) developed the first switch-based method to integrate gene expression data within the FBA framework. In this method, inactive genes were determined based on their non-detection across the microarray replicates for a given condition, and the maximum flux through reactions associated with such inactive genes is set to zero. To overcome pitfalls due to experimental noise, Akesson *et al.* performed a manual assessment of the inactive gene set for false negatives, and re-enabled any reaction flux associated with such manually identified false negatives. Subsequent to the work of Akesson *et al.*, Becker *et al.* (Becker and Palsson, 2008) have developed a switch-based method, Gene inactivity moderated by metabolism and expression (GIMME), which is widely used to build context-specific metabolic models. Unlike Akesson *et al*, GIMME semi-automatically assesses the inactive gene set for potential false negatives, and re-enables any reaction flux associated with such false negatives. In addition, GIMME has the advantage of reporting an inconsistency score between the gene expression data and the predicted flux distribution. Although the methods in this category were initially developed to integrate transcriptomic data, overtime these methods (Becker and Palsson, 2008; Schmidt et al., 2013) have also been utilized to incorporate other types of omics data, such as proteomics and metabolomics, within the FBA framework.

In contrast to the switch-based methods, Colijn *et al.* (Colijn et al., 2009) developed a valve-based method, E-flux, to integrate gene expression data within the FBA framework. In E-flux, bounds are set such that the maximum allowable flux through each reaction is a function of the normalized expression of the associated genes (Fig. 2). Chandrasekaran *et al.* (Chandrasekaran and Price, 2010) developed another valve-based method probabilistic regulation of metabolism (PROM) which integrates both gene expression data and transcriptional regulatory networks (TRNs) within the FBA framework, and thus, this method is discussed in the penultimate section. In summary, switch-based methods lead to a binary classification of flux constraints while valve-based methods (Colijn et al., 2009; Chandrasekaran and Price, 2010; Fang et al., 2012) impose a continuum of flux constraints (Fig. 2). Note that all methods in the first category make use of LP and a biologically relevant objective function to predict flux distribution and generate context-specific metabolic models.

The second category includes methods that integrate gene expression data to build context-specific metabolic models without imposing a biologically relevant objective function. Shlomi *et al.* (Shlomi et al., 2008; Zur et al., 2010) developed the first method in this category, integrated metabolic analysis tool (iMAT), which can integrate transcriptomics and proteomics data to build tissue-specific metabolic models in multicellular organisms. iMAT uses omics data to classify genes as highly, moderately or lowly expressed in a given condition, and subsequently, resorts to mixed integer linear programming (MILP) to generate a context-specific metabolic model where the presence of reactions associated with highly expressed genes is maximized while the presence of reactions associated with lowly expressed genes is minimized. Agren *et al.* (Agren et al., 2012) developed a similar MILP based method, integrative network inference for tissues (INIT), which can also integrate diverse omics data to generate context-specific metabolic models. In contrast to iMAT, INIT enforces the additional constraint of positive production of experimentally determined metabolites based on metabolomics data while building the tissue-specific metabolic models. For normal tissues in mammalian systems (unlike cancerous tumours), it is inappropriate to use growth or proliferation as the relevant objective function. Thus, iMAT and INIT are better suited to generate context-specific metabolic models for normal tissues in comparison to methods in the first category. An extension of INIT, task-driven integrative network inference for tissues (tINIT), has been recently proposed which unlike INIT and iMAT enables the user to also specify a set of required metabolic functions to be satisfied by final context-specific metabolic model (Agren et al., 2014). Thus, tINIT is a MILP based method which also shares principles with the first category of LP based methods.

The third category includes methods that were envisaged to build mammalian tissue-specific metabolic models. These methods integrate diverse omics data and literature based annotations to define two sets of reactions, core and non-core, for a specific tissue type. Reactions in the core set are highly likely to be functional while those in the non-core set are unlikely to be functional in the tissue under consideration. After defining the core and non-core set of reactions for a given tissue, these methods attempt to iteratively remove each non-core reaction subject to preserving the functionality of the core reaction set, and this leads to a tissue-specific metabolic model from a generic metabolic model. Some of the methods which fall into this third category include the model building algorithm (MBA) (Jerby et al., 2010), the metabolic context-specificity assessed by deterministic reaction evaluation (mCADRE) (Wang et al., 2012), fast reconstruction of compact context-specific metabolic network model (FASTCORE) (Vlassis et al., 2014) and cost optimization reaction dependency assessment (CORDA) (Schultz and Qutub, 2016).

We emphasize that some of the proposed methods to integrate omics data within the FBA framework, such as metabolic adjustment by differential expression (MADE) (Jensen and Papin, 2011) and temporal expression-based analysis of metabolites (TEAM) (Collins et al., 2012), cannot be clearly assigned to any of the above mentioned three categories. MADE requires expression data from two or more successive conditions, such as a temporal expression profile, to determine differential expression of genes. Based on statistically significant changes in expression of genes between successive conditions, MADE infers highly and lowly expressed reactions for a sequence of conditions without the need for a predefined threshold. Finally, MADE solves a single MILP problem to generate context-specific metabolic model that best recapitulates the expression dynamics across successive conditions. On the other hand, TEAM is a fusion method based on dynamic FBA (Mahadevan et al., 2002) and GIMME to integrate temporal gene expression data within the FBA framework to predict time course flux distributions. In Supplementary Table 1, we list all the different methods developed so far to integrate omics data within the FBA framework.

## 4. Incorporation of Boolean transcriptional regulatory networks to build integrated regulatory-metabolic models

In the previous section, we discussed several existing methods to integrate multi-omics data as additional flux constraints within the FBA framework to understand the condition-specific regulation of metabolism. Note that such methods are inherently descriptive rather than predictive in nature (Covert et al., 2004; Chandrasekaran and Price, 2010). That is, such methods are only able to describe the regulation of metabolism in conditions with available omics data but cannot predict in conditions lacking omics data. Moreover, such methods do not explicitly incorporate available information on transcriptional regulatory interactions within the FBA framework, and thus, their predictions are restricted to metabolic genes and associated reactions (Covert et al., 2001; Covert and Palsson, 2002; Covert et al., 2004; Herrgard et al., 2006; Shlomi et al., 2007; Goelzer et al., 2008; Samal and Jain, 2008; Chandrasekaran and Price, 2010).

Large-scale transcriptional regulatory networks (TRNs) have been reconstructed for several organisms including *Escherichia coli* (Gama-Castro et al., 2016), *Bacillus subtilis* (Sierro et al., 2008; Nicolas et al., 2012; Kumar et al., 2015), *Saccharomyces cerevisiae* (Teixeira et al., 2014) and humans (Gerstein et al., 2012) based on diverse biological datasets. Our current knowledge of these TRNs is mostly limited to the set of interactions and information on the nature of interactions (activating or repressing) between transcriptional factors (TFs) and their regulated genes. But inadequate information on parameters characterizing most interactions render detailed modelling of large-scale TRNs based on differential equations infeasible (Bornholdt, 2005). Boolean networks (Kauffman, 1969a; Kauffman, 1969b; Thomas, 1973; Kauffman, 1993), an alternate approach first proposed by Stuart Kauffman, has been widely used to study the qualitative dynamics of large-scale TRNs (de Jong, 2002). In the Boolean framework, each gene in the network is in one of two states, active or inactive. The state of each gene at a given time is determined by the state of its regulating gene(s) at the previous time based on a Boolean input function. The state of genes’ in a Boolean network are either updated synchronously or asynchronously in a discrete time setting. Thus, Boolean networks provide a qualitative description of the dynamics of large-scale TRNs (Bornholdt, 2005).

Integrated regulatory-metabolic models (Covert et al., 2004; Herrgard et al., 2006; Goelzer et al., 2008) have been manually reconstructed for some model organisms by unifying Boolean models of TRNs with FBA models of metabolic networks (Fig. 3). Covert *et al.* (Covert et al., 2004) were the first to reconstruct such a genome-scale integrated regulatory-metabolic model iMC1010 for *E. coli*, which was obtained by fusing a Boolean model of the TRN with the genome-scale metabolic network iJR904. iMC1010 accounts for 1010 genes in *E. coli* of which 104 genes code for 103 TFs and 906 genes code for different enzymes in the metabolic network iJR904. Moreover, the activity of genes in iMC1010 is determined by Boolean logical functions that depend on the state of its regulating TFs in the TRN. In addition, the Boolean logical functions that determine the activity of genes in iMC1010 also depend on the presence or absence of certain metabolites in the environment, flux of metabolic reactions and certain stimuli such as heat shock, stress, etc. Since, constraint-based FBA framework assumes steady state to predict the flux distribution in a metabolic network, the method cannot predict internal metabolite concentrations. Thus, in iMC1010, the authors use the flux of internal reactions as surrogate to model the allosteric regulation of proteins which is dependent on internal metabolite concentrations. Similar integrated regulatory-metabolic models have also been manually reconstructed for *B. subtilis* (Goelzer et al., 2008) and S. *cerevisiae* (Herrgard et al., 2006). Note that these integrated regulatory-metabolic models can also predict the metabolic phenotype of TF perturbations.

**Figure 3.**
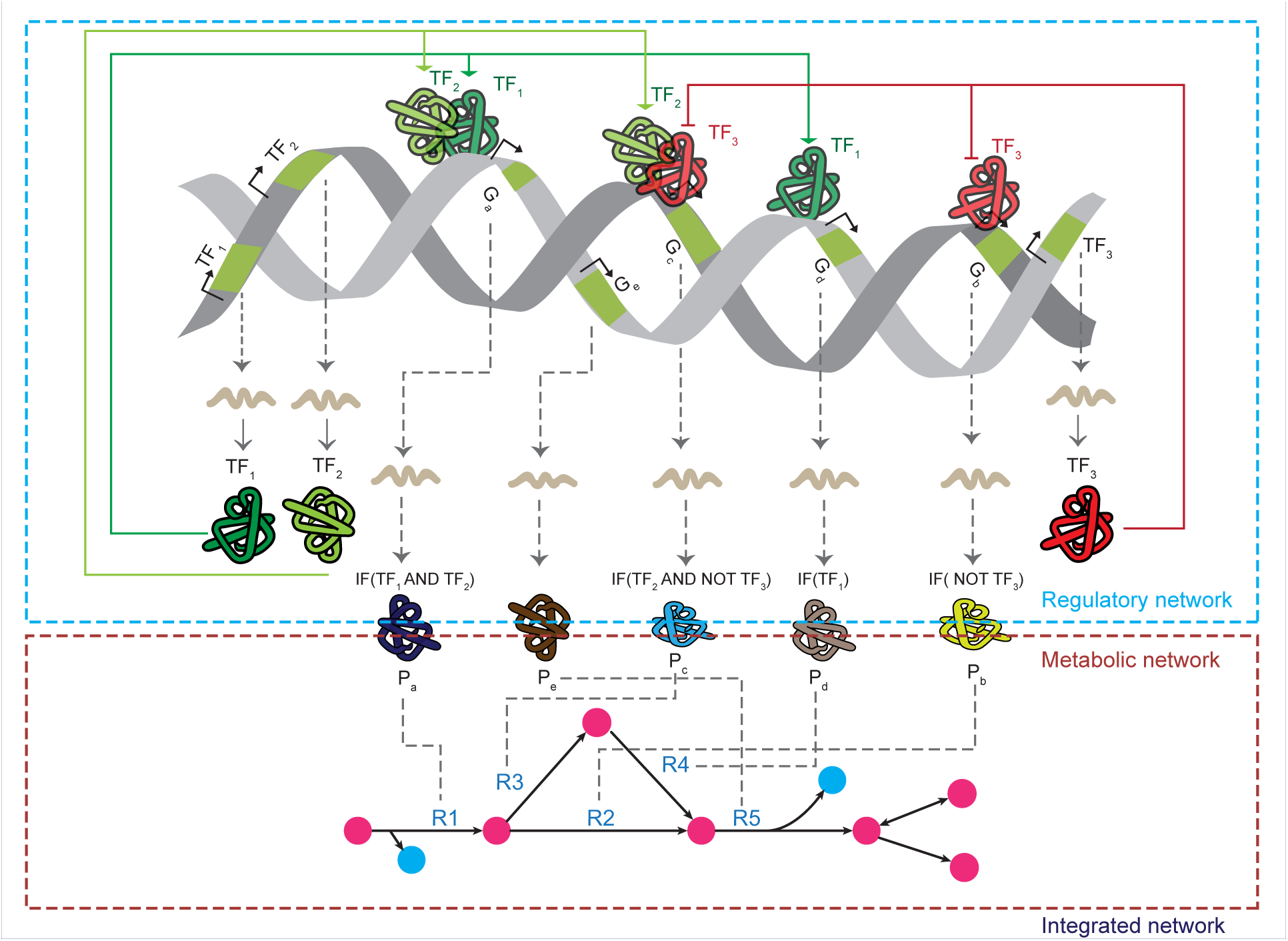
Incorporation of Boolean transcriptional regulatory networks within FBA framework to build integrated regulatory-metabolic models. In this approach, the known TRN of an organism is captured in a Boolean model where the activity of each gene is determined by the state of its regulating genes coding for TFs. For example, gene G_c_ coding for enzyme P_c_ associated with reaction R3 is controlled by transcription factors TF_2_ and TF_3_, where TF_3_ represses the transcription of G_c_ while TF_2_ activates it. Thus, the Boolean rule for gene G_c_ captures this combinatorial regulation by TF_2_ and TF_3_. The figure also shows a representative metabolic network where constituent reactions are directly controlled by the state of genes coding for enzymes and indirectly controlled by genes coding for TFs. Such an integrated regulatory-metabolic model can be investigated using the rFBA (Covert et al., 2001; Covert et al., 2004) approach.

Regulatory flux balance analysis (rFBA) (Covert et al., 2001; Covert et al., 2004) is a method that has been developed to simulate these integrated regulatory-metabolic models incorporating TRNs as Boolean networks. rFBA simulates the growth of an organism in batch cultures to predict a series of steady state flux distributions corresponding to changes in the growth environment in successive time intervals. For each time interval, rFBA determines the state of the regulatory network or activity of genes in the integrated model based on logical functions in the Boolean regulatory network where the present regulatory state is also influenced by the metabolic state in the previous time interval. Next, rFBA uses the predicted activity of genes in the current time interval to constrain the flux of associated reactions within the FBA framework and compute the steady state flux distribution or metabolic state for the current interval (Fig. 3). Note that rFBA also predicts the temporal dynamics of concentration of external substrates available for uptake, concentration of secreted by-products and cell growth. It has been shown that the application of rFBA to the integrated regulatory-metabolic model iMC1010 for *E. coli* can significantly increase the ability to predict knockout phenotypes across diverse environmental conditions (Covert et al., 2004).

One limitation of rFBA is that it arbitrarily chooses one optimal flux distribution as the metabolic state of the integrated model from a space of alternate optimal flux distributions at each time interval, and steady-state regulatory flux balance analysis (SR-FBA) (Shlomi et al., 2007) is a MILP based method which tries to overcome this limitation. Another limitation of using rFBA is that constraint-based methods do not predict the kinetics of internal metabolite concentrations which is important to understand the allosteric regulation of metabolism. On the other hand, detailed kinetic models (Chassagnole et al., 2002; Bettenbrock et al., 2007) can properly account for allosteric regulation but such models are limited to specific metabolic pathways due to paucity of kinetic data. For example, such a detailed kinetic model (Bettenbrock et al., 2007) with less than 50 reactions has been built to study carbon catabolite repression in *E. coli*. Covert *et al.* (Covert et al., 2008) have developed a hybrid framework, integrated FBA (iFBA), which combines rFBA for genome-scale networks and kinetic models for smaller well characterized pathways to improve the predictive ability of integrated models.

## 5. Automated reconstruction of integrated regulatory-metabolic models

In the previous section, we reviewed manually reconstructed integrated regulatory-metabolic models where transcriptional regulatory interactions is represented as Boolean rules. However, manual reconstruction of Boolean models of TRNs from available information on interactions between TFs and their regulated genes can be a slow and tedious process. Firstly, it is extremely difficult to infer a single well-defined set of Boolean logical functions for genes in the TRN based on experimentally determined expression states in different conditions (Henry et al., 2013). Secondly, the Boolean approximation where each gene can have only two states, active or inactive, is an over-simplified view of the system that is inadequate to capture the complex regulation of enzymes in many situations. Thus, Boolean network based integrated regulatory-metabolic models have been manually reconstructed for only three organisms (Covert et al., 2004; Herrgard et al., 2006; Goelzer et al., 2008) to date.

On the other hand, large-scale sequencing projects such as ENCODE and modENCODE (Gerstein et al., 2010; mod et al., 2010; Gerstein et al., 2012) have generated vast amounts of regulatory information for humans and model organisms, mainly, in the form of RNA-seq and ChIP-seq data in diverse biological contexts. Moreover, with the rapid fall in the cost of sequencing, experimental biologists are routinely generating RNA-seq and ChIP-seq data for their pet organisms. RNA-seq data gives information on mRNA abundances or gene expression in a given condition and ChIP-seq data on genome-wide binding profile for TFs or transcriptional regulatory interactions. Combination of ChIP-seq and RNA-seq data can be used to reconstruct the TRN of an organism, and thus, such information can be incorporated within genome-scale metabolic reconstructions to build integrated regulatory-metabolic models (Fig. 4). Given the slow pace of manual reconstruction process, automated approaches combining information on regulatory network architecture and gene expression data are best suited to address the challenge of turning the deluge of high-throughput data into predictive systems biology models.

**Figure 4.**
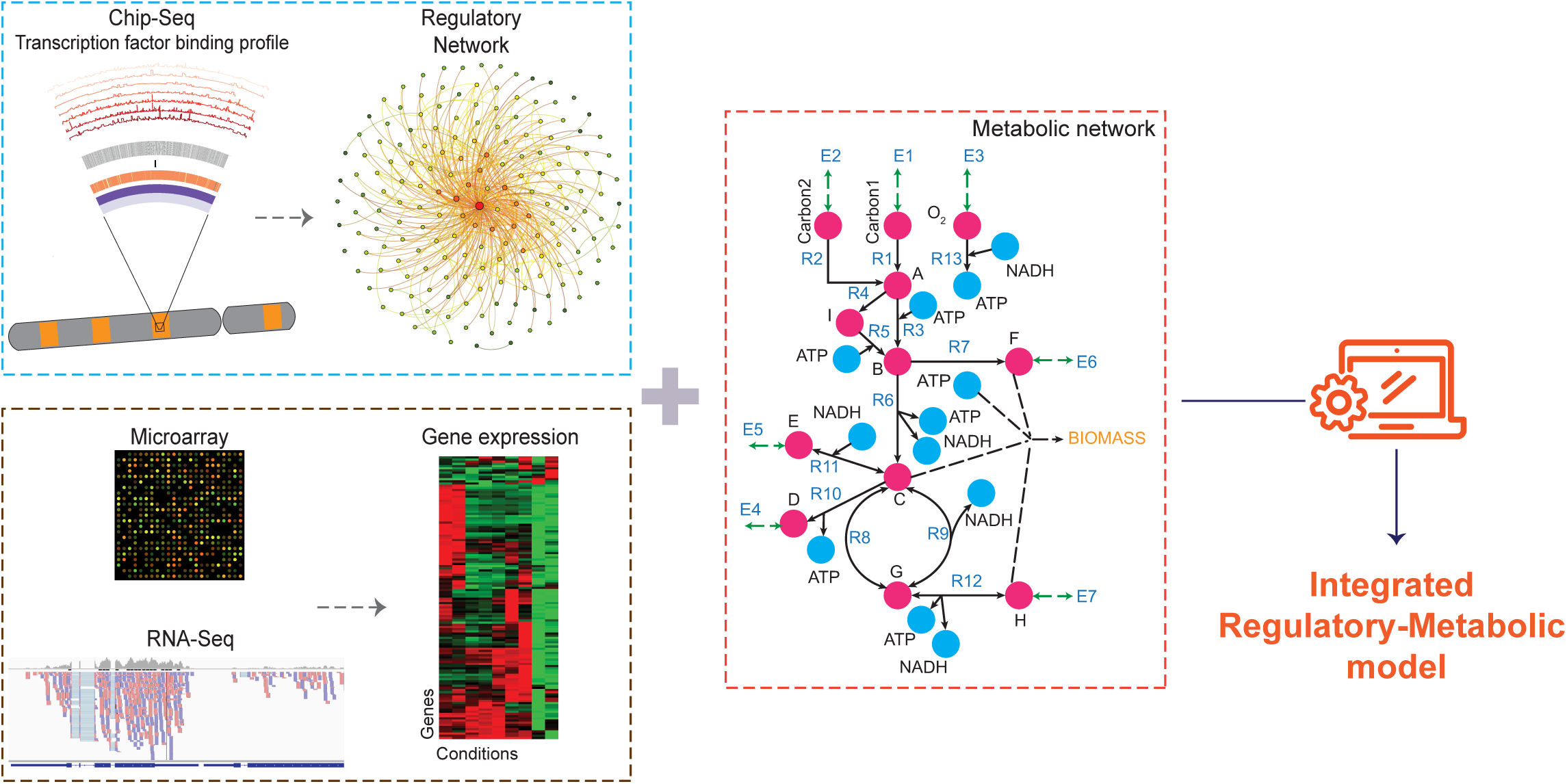
Automated reconstruction of integrated regulatory-metabolic models. Integrated regulatory-metabolic models are holistic and predictive models that capture the complex interactions between transcriptional regulatory network (TRN) and metabolic network of an organism. Such integrated regulatory-metabolic models can be reconstructed based on information on the TRN derived from genome-wide ChiP-seq experiments and gene expression profile from microarray or RNA-seq experiments. Efficient automated algorithms are then employed to integrate diverse regulatory information into the genome-scale metabolic network to generate high quality integrated regulatory-metabolic models.

In this direction, probabilistic regulation of metabolism (PROM) (Chandrasekaran and Price, 2010), is the only automated method to date. PROM combines information on TRN architecture and gene expression data within the FBA framework to build an integrated model. In PROM, the set of interactions between TFs and target genes in the TRN is used along with gene expression data across multiple conditions to predict the likelihood of the expression of a target gene given the expression state of the controlling TF. In PROM, such probabilities are then used to constrain the maximum allowable flux through reactions associated with the target genes before computing the flux distribution using FBA. Thus, PROM can predict the consequences of TF perturbations unlike methods in the first approach. Also, the prediction accuracy achieved by PROM was similar to the manually reconstructed integrated regulatory-metabolic model of *E. coli*.

Nevertheless, the existing methods including PROM have limitations that provide scope for future improvements. Firstly, the existing automated method does not make use of information on environment-dependent or condition-specific regulation. Secondly, the existing methods are designed to integrate high-confidence regulatory interactions from manually reconstructed TRNs. In contrast, regulatory interactions inferred from ChIP-seq data are likely to be very noisy, and future methods should address this challenge. Thirdly, existing methods do not account for available information on feedback regulation of enzymes by intracellular metabolites. Addressing this challenge may require transition to hybrid models where a combination of constraint-based and kinetic approaches will be necessary to simulate metabolic networks (Covert et al., 2008). Finally, the existing automated method to build integrated regulatory-metabolic model is static in time and not designed to incorporate temporal expression data. We expect future automated methods will overcome some of the above-mentioned limitations.

## 6. Conclusion

In this review, we have presented an overview of three different approaches for integrating available regulatory information within constraint-based FBA models of genome-scale metabolic reconstructions. Previous reviews (Blazier and Papin, 2012; Hyduke et al., 2013; Estévez and Nikoloski, 2014) in this area have largely focused on the first approach where considerable research has led to several methods where omics data is directly integrated within the FBA framework as additional flux constraints (Supplementary Table 1). Such methods including GIMME (Becker and Palsson, 2008), iMAT (Shlomi et al., 2008), INIT (Agren et al., 2012) and MBA (Jerby et al., 2010) have been successful in building context-and tissue-specific metabolic models. Notably, the methods in the first approach are rather descriptive than predictive as they do not explicitly incorporate the available information on regulatory interactions in the TRN (Chandrasekaran and Price, 2010). In this review, we also focus on two other approaches to build integrated regulatory-metabolic models. First such approach involves manual reconstruction of integrated regulatory-metabolic models where transcriptional regulatory information is represented as Boolean networks (Covert et al., 2004; Herrgard et al., 2006; Goelzer et al., 2008). Such Boolean based integrated regulatory-metabolic models can be studied using rFBA and allied approaches (Shlomi et al., 2007; Samal and Jain, 2008). Since manual reconstruction of integrated models is a painstaking and time consuming process, the need of the hour is to develop automated methods to incorporate the flood of next generation sequencing data such as RNA-seq and ChIP-seq within genome-scale metabolic reconstructions to build integrated regulatory-metabolic models. Such automated methods will eventually enable the systems biology community to reconstruct whole-scale models (Karr et al., 2012) for several organisms.

## Acknowledgments

We would like to apologize to the authors of other relevant articles whose work was not cited in this review due to limited space. AS acknowledges support from Department of Science and Technology (DST) India start-up grant (YSS/2015/000060), Ramanujan fellowship (SB/S2/RJN-006/2014), Max Planck India mobility grant and IMSc PRISM project (XII Plan).

### Supplementary Material

**Supplementary Table 1:** Comprehensive list of methods proposed so far to integrate omics data within constraint-based metabolic models. Note that some of the methods listed here cannot be clearly assigned to any of the three categories discussed in section 3 of the main text.

## References

Agren, R., Bordel, S., Mardinoglu, A., Pornputtapong, N., Nookaew, I., Nielsen, J., 2012. Reconstruction of genome-scale active metabolic networks for 69 human cell types and 16 cancer types using INIT. PLoS Comput Biol 8, e1002518.

Agren, R., Liu, L, Shoaie, S., Vongsangnak, W., Nookaew, I., Nielsen, J., 2013. The RAVEN toolbox and its use for generating a genome-scale metabolic model for Penicillium chrysogenum. PLoS Comput Biol 9, e1002980.

Agren, R., Mardinoglu, A., Asplund, A., Kampf, C, Uhlen, M., Nielsen, J., 2014. Identification of anticancer drugs for hepatocellular carcinoma through personalized genome-scale metabolic modeling. Mol Syst Biol 10, 721.

Åkesson, M., Förster, J., Nielsen, J., 2004. Integration of gene expression data into genome-scale metabolic models. Metabolic engineering 6, 285–293.

Beard, D.A., Liang, S.D., Qian, H., 2002. Energy balance for analysis of complex metabolic networks. Biophys J 83, 79–86.

Becker, S.A., Palsson, B.O., 2008. Context-specific metabolic networks are consistent with experiments. PLoS Comput Biol 4, e1000082.

Bettenbrock, K., Sauter, T., Jahreis, K., Kremling, A., Lengeler, J.W., Gilles, E.D., 2007. Correlation between growth rates, EIIACrr phosphorylation, and intracellular cyclic AMP levels in Escherichia coli K-12. J Bacteriol 189, 6891–6900.

Blazier, A.S., Papin, J.A., 2012. Integration of expression data in genome-scale metabolic network reconstructions. Front Physiol 3.

Bordbar, A., Mo, M.L., Nakayasu, E.S., Schrimpe-Rutledge, A.C., Kim, Y.M., Metz, T.O., Jones, M.B., Frank, B.C., Smith, R.D., Peterson, S.N., 2012. Model-driven multi-omic data analysis elucidates metabolic immunomodulators of macrophage activation. Molecular systems biology 8, 558.

Bornholdt, S., 2005. Systems biology. Less is more in modeling large genetic networks. Science 310, 449–451.

Burgard, A.P., Pharkya, P., Maranas, C.D., 2003. Optknock: a bilevel programming framework for identifying gene knockout strategies for microbial strain optimization. Biotechnology and bioengineering 84, 647–657.

Chandrasekaran, S., Price, N.D., 2010. Probabilistic integrative modeling of genome-scale metabolic and regulatory networks in Escherichia coli and Mycobacterium tuberculosis. Proceedings of the National Academy of Sciences 107, 17845–17850.

Chang, A., Scheer, M., Grote, A., Schomburg, I., Schomburg, D., 2009. BRENDA, AMENDA and FRENDA the enzyme information system: new content and tools in 2009. Nucleic Acids Res 37, D588–592.

Chassagnole, C., Noisommit-Rizzi, N., Schmid, J.W., Mauch, K., Reuss, M., 2002. Dynamic modeling of the central carbon metabolism of Escherichia coli. Biotechnology and bioengineering 79, 53–73.

Colijn, C., Brandes, A., Zucker, J., Lun, D.S., Weiner, B., Farhat, M.R., Cheng, T.-Y., Moody, D.B., Murray, M., Galagan, J.E., 2009. Interpreting expression data with metabolic flux models: predicting Mycobacterium tuberculosis mycolic acid production. PLoS Comput Biol 5, e1000489.

Collins, S.B., Reznik, E., Segrè, D., 2012. Temporal expression-based analysis of metabolism. PLoS Comput Biol 8, e1002781.

Covert, M.W., Knight, E.M., Reed, J.L., Herrgard, M.J., Palsson, B.O., 2004. Integrating high-throughput and computational data elucidates bacterial networks. Nature 429, 92–96.

Covert, M.W., Palsson, B.O., 2002. Transcriptional regulation in constraints-based metabolic models of Escherichia coli. J Biol Chem 277, 28058–28064.

Covert, M.W., Palsson, B.O., 2003. Constraints-based models: regulation of gene expression reduces the steady-state solution space. J Theor Biol 221, 309–325.

Covert, M.W., Schilling, C.H., Palsson, B., 2001. Regulation of gene expression in flux balance models of metabolism. J Theor Biol 213, 73–88.

Covert, M.W., Xiao, N., Chen, T.J., Karr, J.R., 2008. Integrating metabolic, transcriptional regulatory and signal transduction models in Escherichia coli. Bioinformatics 24, 2044–2050.

de Jong, H., 2002. Modeling and simulation of genetic regulatory systems: a literature review. J Comput Biol 9, 67–103.

Duarte, N.C., Becker, S.A., Jamshidi, N., Thiele, I., Mo, M.L., Vo, T.D., Srivas, R., Palsson, B.O., 2007. Global reconstruction of the human metabolic network based on genomic and bibliomic data. Proc Natl Acad Sci U S A 104, 1777–1782.

Durot, M., Bourguignon, P.-Y., Schachter, V., 2009. Genome-scale models of bacterial metabolism: reconstruction and applications. FEMS microbiology reviews 33, 164–190.

Edwards, J.S., Ibarra, R.U., Palsson, B.O., 2001. In silico predictions of Escherichia coli metabolic capabilities are consistent with experimental data. Nat Biotechnol 19, 125–130.

Estévez, S.R., Nikoloski, Z., 2014. Generalized framework for context-specific metabolic model extraction methods. Front. Plant Sci 5, 10.3389.

Fang, X., Wallqvist, A., Reifman, J., 2012. Modeling phenotypic metabolic adaptations of Mycobacterium tuberculosis H37Rv under hypoxia. PLoS Comput Biol 8, e1002688.

Feist, A.M., Herrgård, M.J., Thiele, I., Reed, J.L., Palsson, B.O., 2009. Reconstruction of biochemical networks in microorganisms. Nature Reviews Microbiology 7, 129–143.

Feist, A.M., Palsson, B.O., 2008. The growing scope of applications of genome-scale metabolic reconstructions using Escherichia coli. Nat Biotechnol 26, 659–667.

Feist, A.M., Palsson, B.O., 2010. The biomass objective function. Curr Opin Microbiol 13, 344–349.

Fell, D.A., Small, J.R., 1986. Fat synthesis in adipose tissue. An examination of stoichiometric constraints. Biochem J 238, 781–786.

Fleischmann, R.D., Adams, M.D., White, O., Clayton, R.A., Kirkness, E.F., Kerlavage, A.R., Bult, C.J., Tomb, J.F., Dougherty, B.A., Merrick, J.M., et al., 1995. Whole-genome random sequencing and assembly of Haemophilus influenzae Rd. Science 269, 496–512.

Folger, O., Jerby, L, Frezza, C., Gottlieb, E., Ruppin, E., Shlomi, T., 2011. Predicting selective drug targets in cancer through metabolic networks. Mol Syst Biol 7, 501.

Forster, J., Famili, I., Fu, P., Palsson, B.O., Nielsen, J., 2003. Genome-scale reconstruction of the Saccharomyces cerevisiae metabolic network. Genome Res 13, 244–253.

Gama-Castro, S., Salgado, H., Santos-Zavaleta, A., Ledezma-Tejeida, D., Muniz-Rascado, L, Garcia-Sotelo, J.S., Alquicira-Hernandez, K., Martinez-Flores, I., Pannier, L., Castro-Mondragon, J.A., Medina-Rivera, A., Solano-Lira, H., Bonavides-Martinez, C., Perez-Rueda, E., Alquicira-Hernandez, S., Porron-Sotelo, L., Lopez-Fuentes, A., Hernandez-Koutoucheva, A., Moral-Chavez, V.D., Rinaldi, F., Collado-Vides, J., 2016. RegulonDB version 9.0: high-level integration of gene regulation, coexpression, motif clustering and beyond. Nucleic Acids Res 44, D133–143.

Gerstein, M.B., Kundaje, A., Hariharan, M., Landt, S.G., Yan, K.K., Cheng, C., Mu, X.J., Khurana, E., Rozowsky, J., Alexander, R., Min, R., Alves, P., Abyzov, A., Addleman, N., Bhardwaj, N., Boyle, A.P., Cayting, P., Charos, A., Chen, D.Z., Cheng, Y., Clarke, D., Eastman, C., Euskirchen, G., Frietze, S., Fu, Y., Gertz, J., Grubert, F., Harmanci, A., Jain, P., Kasowski, M., Lacroute, P., Leng, J., Lian, J., Monahan, H., O’Geen, H., Ouyang, Z., Partridge, E.C., Patacsil, D., Pauli, F., Raha, D., Ramirez, L., Reddy, T.E., Reed, B., Shi, M., Slifer, T., Wang, J., Wu, L., Yang, X., Yip, K.Y., Zilberman-Schapira, G., Batzoglou, S., Sidow, A., Farnham, P.J., Myers, R.M., Weissman, S.M., Snyder, M., 2012. Architecture of the human regulatory network derived from ENCODE data. Nature 489, 91–100.

Gerstein, M.B., Lu, Z.J., Van Nostrand, E.L., Cheng, C., Arshinoff, B.I., Liu, T., Yip, K.Y., Robilotto, R., Rechtsteiner, A., Ikegami, K., Alves, P., Chateigner, A., Perry, M., Morris, M., Auerbach, R.K., Feng, X., Leng, J., Vielle, A., Niu, W., Rhrissorrakrai, K., Agarwal, A., Alexander, R.P., Barber, G., Brdlik, C.M., Brennan, J., Brouillet, J.J., Carr, A., Cheung, M.S., Clawson, H., Contrino, S., Dannenberg, L.O., Dernburg, A.F., Desai, A., Dick, L., Dose, A.C., Du, J., Egelhofer, T., Ercan, S., Euskirchen, G., Ewing, B., Feingold, E.A., Gassmann, R., Good, P.J., Green, P., Gullier, F., Gutwein, M., Guyer, M.S., Habegger, L., Han, T., Henikoff, J.G., Henz, S.R., Hinrichs, A., Holster, H., Hyman, T., Iniguez, A.L., Janette, J., Jensen, M., Kato, M., Kent, W.J., Kephart, E., Khivansara, V., Khurana, E., Kim, J.K., Kolasinska-Zwierz, P., Lai, E.C., Latorre, I., Leahey, A., Lewis, S., Lloyd, P., Lochovsky, L., Lowdon, R.F., Lubling, Y., Lyne, R., MacCoss, M., Mackowiak, S.D., Mangone, M., McKay, S., Mecenas, D., Merrihew, G., Miller, D.M., 3rd, Muroyama, A., Murray, J.I., Ooi, S.L., Pham, H., Phippen, T., Preston, E.A., Rajewsky, N., Ratsch, G., Rosenbaum, H., Rozowsky, J., Rutherford, K., Ruzanov, P., Sarov, M., Sasidharan, R., Sboner, A., Scheid, P., Segal, E., Shin, H., Shou, C., Slack, F.J., Slightam, C., Smith, R., Spencer, W.C., Stinson, E.O., Taing, S., Takasaki, T., Vafeados, D., Voronina, K., Wang, G., Washington, N.L., Whittle, C.M., Wu, B., Yan, K.K., Zeller, G., Zha, Z., Zhong, M., Zhou, X., mod, E.C., Ahringer, J., Strome, S., Gunsalus, K.C., Micklem, G., Liu, X.S., Reinke, V., Kim, S.K., Hillier, L.W., Henikoff, S., Piano, F., Snyder, M., Stein, L., Lieb, J.D., Waterston, R.H., 2010. Integrative analysis of the Caenorhabditis elegans genome by the modENCODE project. Science 330, 1775–1787.

Gianchandani, E.P., Chavali, A.K., Papin, J.A., 2010. The application of flux balance analysis in systems biology. Wiley interdisciplinary reviews. Systems biology and medicine 2, 372–382.

Goelzer, A., Bekkal Brikci, F., Martin-Verstraete, I., Noirot, P., Bessieres, P., Aymerich, S., Fromion, V., 2008. Reconstruction and analysis of the genetic and metabolic regulatory networks of the central metabolism of Bacillus subtilis. BMC Syst Biol 2, 20.

Gygi, S.P., Rochon, Y., Franza, B.R., Aebersold, R., 1999. Correlation between protein and mRNA abundance in yeast. Mol Cell Biol 19, 1720–1730.

Heinrich, R., Schuster, S., 1996. The Regulation of Cellular Systems. Springer US, Boston, MA, p. 1 online resource (416 pages).

Henry, A., Moneger, F., Samal, A., Martin, O.C., 2013. Network function shapes network structure: the case of the Arabidopsis flower organ specification genetic network. Mol Biosyst 9, 1726–1735.

Henry, C.S., DeJongh, M., Best, A.A., Frybarger, P.M., Linsay, B., Stevens, R.L., 2010. High-throughput generation, optimization and analysis of genome-scale metabolic models. Nat Biotechnol 28, 977–982.

Herrgard, M.J., Lee, B.S., Portnoy, V., Palsson, B.O., 2006. Integrated analysis of regulatory and metabolic networks reveals novel regulatory mechanisms in Saccharomyces cerevisiae. Genome Res 16, 627–635.

Hyduke, D.R., Lewis, N.E., Palsson, B.O., 2013. Analysis of omics data with genome-scale models of metabolism. Mol Biosyst 9, 167–174.

Imam, S., Schauble, S., Brooks, A.N., Baliga, N.S., Price, N.D., 2015. Data-driven integration of genome-scale regulatory and metabolic network models. Front Microbiol 6, 409.

Jensen, P.A., Papin, J.A., 2011. Functional integration of a metabolic network model and expression data without arbitrary thresholding. Bioinformatics 27, 541–547.

Jerby, L., Shlomi, T., Ruppin, E., 2010. Computational reconstruction of tissue-specific metabolic models: application to human liver metabolism. Mol Syst Biol 6, 401.

Joyce, A.R., Palsson, B.O., 2006. The model organism as a system: integrating ‘omics’ data sets. Nat Rev Mol Cell Biol 7, 198–210.

Kanehisa, M., Goto, S., 2000. KEGG: kyoto encyclopedia of genes and genomes. Nucleic Acids Res 28, 27–30.

Karr, J.R., Sanghvi, J.C., Macklin, D.N., Gutschow, M.V., Jacobs, J.M., Bolival, B., Jr., Assad-Garcia, N., Glass, J.I., Covert, M.W., 2012. A whole-cell computational model predicts phenotype from genotype. Cell 150, 389–401.

Kauffman, K.J., Prakash, P., Edwards, J.S., 2003. Advances in flux balance analysis. Curr Opin Biotechnol 14, 491–496.

Kauffman, S., 1969a. Homeostasis and differentiation in random genetic control networks. Nature 224, 177–178.

Kauffman, S.A., 1969b. Metabolic stability and epigenesis in randomly constructed genetic nets. J Theor Biol 22, 437–467.

Kauffman, S.A., 1993. The origins of order: self-organization and selection in evolution. Oxford University Press, New York.

Kim, J., Reed, J.L., 2014. Refining metabolic models and accounting for regulatory effects. Current opinion in biotechnology 29, 34–38.

Kitano, H., 2002. Computational systems biology. Nature 420, 206–210.

Kumar, S., Vendruscolo, M., Singh, A., Kumar, D., Samal, A., 2015. Analysis of the hierarchical structure of the B. subtilis transcriptional regulatory network. Mol Biosyst 11, 930–941.

Lewis, N.E., Nagarajan, H., Palsson, B.O., 2012. Constraining the metabolic genotype-phenotype relationship using a phylogeny of in silico methods. Nature Reviews Microbiology 10, 291–305.

Machado, D., Herrgård, M., 2014. Systematic evaluation of methods for integration of transcriptomic data into constraint-based models of metabolism. PLoS Comput Biol 10, e1003580.

Mahadevan, R., Edwards, J.S., Doyle, F.J., 2002. Dynamic flux balance analysis of diauxic growth in Escherichia coli. Biophysical journal 83, 1331–1340.

mod, E.C., Roy, S., Ernst, J., Kharchenko, P.V., Kheradpour, P., Negre, N., Eaton, M.L., Landolin, J.M., Bristow, C.A., Ma, L., Lin, M.F., Washietl, S., Arshinoff, B.I., Ay, F., Meyer, P.E., Robine, N., Washington, N.L., Di Stefano, L., Berezikov, E., Brown, C.D., Candeias, R., Carlson, J.W., Carr, A., Jungreis, I., Marbach, D., Sealfon, R., Tolstorukov, M.Y., Will, S., Alekseyenko, A.A., Artieri, C., Booth, B.W., Brooks, A.N., Dai, Q., Davis, C.A., Duff, M.O., Feng, X., Gorchakov, A.A., Gu, T., Henikoff, J.G., Kapranov, P., Li, R., MacAlpine, H.K., Malone, J., Minoda, A., Nordman, J., Okamura, K., Perry, M., Powell, S.K., Riddle, N.C., Sakai, A., Samsonova, A., Sandler, J.E., Schwartz, Y.B., Sher, N., Spokony, R., Sturgill, D., van Baren, M., Wan, K.H., Yang, L., Yu, C., Feingold, E., Good, P., Guyer, M., Lowdon, R., Ahmad, K., Andrews, J., Berger, B., Brenner, S.E., Brent, M.R., Cherbas, L., Elgin, S.C., Gingeras, T.R., Grossman, R., Hoskins, R.A., Kaufman, T.C., Kent, W., Kuroda, M.I., Orr-Weaver, T., Perrimon, N., Pirrotta, V., Posakony, J.W., Ren, B., Russell, S., Cherbas, P., Graveley, B.R., Lewis, S., Micklem, G., Oliver, B., Park, P.J., Celniker, S.E., Henikoff, S., Karpen, G.H., Lai, E.C., MacAlpine, D.M., Stein, L.D., White, K.P., Kellis, M., 2010. Identification of functional elements and regulatory circuits by Drosophila modENCODE. Science 330, 1787–1797.

Nicolas, P., Mader, U., Dervyn, E., Rochat, T., Leduc, A., Pigeonneau, N., Bidnenko, E., Marchadier, E., Hoebeke, M., Aymerich, S., Becher, D., Bisicchia, P., Botella, E., Delumeau, O., Doherty, G., Denham, E.L., Fogg, M.J., Fromion, V., Goelzer, A., Hansen, A., Hartig, E., Harwood, C.R., Homuth, G., Jarmer, H., Jules, M., Klipp, E., Le Chat, L., Lecointe, F., Lewis, P., Liebermeister, W., March, A., Mars, R.A., Nannapaneni, P., Noone, D., Pohl, S., Rinn, B., Rugheimer, F., Sappa, P.K., Samson, F., Schaffer, M., Schwikowski, B., Steil, L., Stulke, J., Wiegert, T., Devine, K.M., Wilkinson, A.J., van Dijl, J.M., Hecker, M., Volker, U., Bessieres, P., Noirot, P., 2012. Condition-dependent transcriptome reveals high-level regulatory architecture in Bacillus subtilis. Science 335, 1103–1106.

Oberhardt, M.A., Palsson, B.O., Papin, J.A., 2009. Applications of genome-scale metabolic reconstructions. Mol Syst Biol 5, 320.

Orth, J.D., Thiele, I., Palsson, B.Ø., 2010. What is flux balance analysis? Nature biotechnology 28, 245–248.

Pal, C., Papp, B., Lercher, M.J., Csermely, P., Oliver, S.G., Hurst, L.D., 2006. Chance and necessity in the evolution of minimal metabolic networks. Nature 440, 667–670.

Palsson, B., 2006. Systems biology: properties of reconstructed networks. Cambridge University Press.

Papp, B., Teusink, B., Notebaart, R.A., 2009. A critical view of metabolic network adaptations. HFSP J 3, 24–35.

Price, N.D., Reed, J.L., Palsson, B.O., 2004. Genome-scale models of microbial cells: evaluating the consequences of constraints. Nat Rev Microbiol 2, 886–897.

Quackenbush, J., 2004. Data standards for ‘omic’ science. Nat Biotechnol 22, 613–614.

Saha, R., Chowdhury, A., Maranas, C.D., 2014. Recent advances in the reconstruction of metabolic models and integration of omics data. Curr Opin Biotechnol 29, 39–45.

Samal, A., Jain, S., 2008. The regulatory network of E. coli metabolism as a Boolean dynamical system exhibits both homeostasis and flexibility of response. BMC Syst Biol 2, 21.

Schellenberger, J., Que, R., Fleming, R.M., Thiele, I., Orth, J.D., Feist, A.M., Zielinski, D.C., Bordbar, A., Lewis, N.E., Rahmanian, S., 2011. Quantitative prediction of cellular metabolism with constraint-based models: the COBRA Toolbox v2. 0. Nature protocols 6, 1290–1307.

Schilling, C.H., Schuster, S., Palsson, B.O., Heinrich, R., 1999. Metabolic pathway analysis: basic concepts and scientific applications in the post-genomic era. Biotechnol Prog 15, 296–303.

Schmidt, B.J., Ebrahim, A., Metz, T.O., Adkins, J.N., Palsson, B.Ø., Hyduke, D.R., 2013. GIM3E: condition-specific models of cellular metabolism developed from metabolomics and expression data. Bioinformatics 29, 2900–2908.

Schuetz, R., Kuepfer, L., Sauer, U., 2007. Systematic evaluation of objective functions for predicting intracellular fluxes in Escherichia coli. Mol Syst Biol 3, 119.

Schultz, A., Qutub, A.A., 2016. Reconstruction of Tissue-Specific Metabolic Networks Using CORDA. PLoS Comput Biol 12, e1004808.

Shlomi, T., Cabili, M.N., Herrgård, M.J., Palsson, B.Ø., Ruppin, E., 2008. Network-based prediction of human tissue-specific metabolism. Nature biotechnology 26, 1003–1010.

Shlomi, T., Eisenberg, Y., Sharan, R., Ruppin, E., 2007. A genome-scale computational study of the interplay between transcriptional regulation and metabolism. Mol Syst Biol 3, 101.

Sierro, N., Makita, Y., de Hoon, M., Nakai, K., 2008. DBTBS: a database of transcriptional regulation in Bacillus subtilis containing upstream intergenic conservation information. Nucleic Acids Res 36, D93–96.

Szappanos, B., Kovacs, K., Szamecz, B., Honti, F., Costanzo, M., Baryshnikova, A., Gelius-Dietrich, G., Lercher, M.J., Jelasity, M., Myers, C.L., Andrews, B.J., Boone, C., Oliver, S.G., Pal, C., Papp, B., 2011. An integrated approach to characterize genetic interaction networks in yeast metabolism. Nat Genet 43, 656–662.

Teixeira, M.C., Monteiro, P.T., Guerreiro, J.F., Goncalves, J.P., Mira, N.P., dos Santos, S.C., Cabrito, T.R., Palma, M., Costa, C., Francisco, A.P., Madeira, S.C., Oliveira, A.L., Freitas, AT., Sa-Correia, I., 2014. The YEASTRACT database: an upgraded information system for the analysis of gene and genomic transcription regulation in Saccharomyces cerevisiae. Nucleic Acids Res 42, D161–166.

ter Kuile, B.H., Westerhoff, H.V., 2001. Transcriptome meets metabolome: hierarchical and metabolic regulation of the glycolytic pathway. FEBS Lett 500, 169–171.

Thiele, I., Palsson, B.O., 2010. A protocol for generating a high-quality genome-scale metabolic reconstruction. Nat Protoc 5, 93–121.

Thiele, I., Swainston, N., Fleming, R.M., Hoppe, A., Sahoo, S., Aurich, M.K., Haraldsdottir, H., Mo, M.L., Rolfsson, O., Stobbe, M.D., Thorleifsson, S.G., Agren, R., Bolling, C., Bordel, S., Chavali, A.K., Dobson, P., Dunn, W.B., Endler, L., Hala, D., Hucka, M., Hull, D., Jameson, D., Jamshidi, N., Jonsson, J.J., Juty, N., Keating, S., Nookaew, I., Le Novere, N., Malys, N., Mazein, A., Papin, J.A., Price, N.D., Selkov, E., Sr., Sigurdsson, M.I., Simeonidis, E., Sonnenschein, N., Smallbone, K., Sorokin, A., van Beek, J.H., Weichart, D., Goryanin, I., Nielsen, J., Westerhoff, H.V., Kell, D.B., Mendes, P., Palsson, B.O., 2013. A community-driven global reconstruction of human metabolism. Nat Biotechnol 31, 419–425.

Thomas, R., 1973. Boolean formalization of genetic control circuits. Journal of theoretical biology 42, 563–585.

Tomb, J.F., White, O., Kerlavage, A.R., Clayton, R.A., Sutton, G.G., Fleischmann, R.D., Ketchum, K.A., Klenk, H.P., Gill, S., Dougherty, B.A., Nelson, K., Quackenbush, J., Zhou, L., Kirkness, E.F., Peterson, S., Loftus, B., Richardson, D., Dodson, R., Khalak, H.G., Glodek, A., McKenney, K., Fitzegerald, L.M., Lee, N., Adams, M.D., Hickey, E.K., Berg, D.E., Gocayne, J.D., Utterback, T.R., Peterson, J.D., Kelley, J.M., Cotton, M.D., Weidman, J.M., Fujii, C., Bowman, C., Watthey, L., Wallin, E., Hayes, W.S., Borodovsky, M., Karp, P.D., Smith, H.O., Fraser, C.M., Venter, J.C., 1997. The complete genome sequence of the gastric pathogen Helicobacter pylori. Nature 388, 539–547.

Varma, A., Palsson, B.O., 1994. Metabolic Flux Balancing: Basic Concepts, Scientific and Practical Use. Nat Biotech 12, 994–998.

Vlassis, N., Pacheco, M.P., Sauter, T., 2014. Fast reconstruction of compact context-specific metabolic network models. PLoS Comput Biol 10, e1003424.

Wang, Y., Eddy, J.A., Price, N.D., 2012. Reconstruction of genome-scale metabolic models for 126 human tissues using mCADRE. BMC systems biology 6, 1.

Watson, M., 1984. Metabolic maps for the Apple II. Biochemical Society Transactions 12, 1093–1094.

Yang, C., Hua, Q., Shimizu, K., 2002. Integration of the information from gene expression and metabolic fluxes for the analysis of the regulatory mechanisms in Synechocystis. Appl Microbiol Biotechnol 58, 813–822.

Zur, H., Ruppin, E., Shlomi, T., 2010. iMAT: an integrative metabolic analysis tool. Bioinformatics 26, 3140–3142.

